# A densely sampled and richly annotated acoustic dataset from a wild bird population

**DOI:** 10.1101/2023.07.03.547484

**Authors:** Nilo Merino Recalde, Andrea Estandía, Loanne Pichot, Antoine Vansse, Ella F. Cole, Ben C. Sheldon

## Abstract

We present a high-resolution, densely-sampled dataset of wild bird songs collected over multiple years from a single population of Great Tits (*Parus major*) in the UK. The dataset includes over 1,100,000 individual acoustic units from 109,963 richly annotated songs, sung by more than 400 individual birds, and provides unprecedented detail on the vocal behaviour of wild birds. Here, we describe the data collection and processing procedures and provide a summary of the data. We also discuss potential research questions that can be addressed using this dataset, including behavioural repeatability and stability, links between vocal performance and reproductive success, the timing of song production, syntactic organisation of song production, and song learning in the wild. We have made the dataset and associated software tools publicly available with the aim that other researchers can benefit from this resource and use it to further our understanding of bird vocal behaviour in the wild.

## Introduction & background

Despite a long history of scientific interest from disciplines as diverse as behavioural ecology, neurobiology and physiology, there is still much to learn regarding the evolution and function of animal vocalisations. Ongoing research covers a wide range of topics, including speech recognition and language evolution in humans, animal welfare, and even fish vocal communication.

Studying animal vocalisations provides us with valuable insights into the dynamics of social interactions and reproductive strategies. Vocalisations often convey essential information about individual state and identity (Lehmann and Seufert 2017; Linhart et al. 2019), group cohesion, and social structure (Bell et al. 2010; Engesser and Manser 2022; Radford and Ridley 2007), as well as playing a significant role in shaping social relationships, mate selection, and parental care (Behr and von Helversen 2004; Gerhardt 1991; Pitcher et al. 2010; Roulin 2001).

Animal vocalisations, particularly those of birds, have long been a focus of research for those interested in social learning and cultural evolution. This interest dates back at least to the pioneering work of Marler and Thorpe with Chaffinches and White-crowned Sparrows (Marler and Tamura 1964, 1962; Marler 1952; Thorpe 1958), which paved the way for what continues to be a thriving field today (see Mets and Brainard 2019; Riebel et al. 2015; Williams and Lachlan 2021; Youngblood and Lahti 2022).

In addition, and from a more mechanistic point of view, they offer a window into the physiological and neural mechanisms underlying vocal production and perception, as well as the consolidation of memories and motor coordination, to name but a few (Davenport and Jarvis 2023).

Beyond their fundamental scientific importance, animal vocalisations have practical applications in various fields. For example, there is increasing recognition of their potential as a non-invasive tool for monitoring populations. By analysing entire soundscapes, researchers can gather crucial information about population dynamics, species distribution, and the presence of rare or elusive species (Kahl et al. 2021; Sethi et al. 2020; Sugai et al. 2019).

However, despite the growing interest in animal vocalisations and their potential applications, publicly available data from wild populations are still scarce—with the notable exception of projects like xeno-canto. This can severely limit researchers’ ability to ask questions that require large datasets to answer, such as those about social learning, vocal development, large-scale cultural diversity, and the syntactic structure of animal vocalisations (Aplin 2019; Kollmorgen et al. 2020; Lachlan et al. 2018; Sainburg et al. 2019).

Indeed, while controlled laboratory settings allow researchers to track vocal development and production in minute detail, it is much harder to obtain finely-grained data from animals in their natural habitats.

The process of collecting such data can be quite demanding and requires significant time, technical expertise, and resources: this includes both data collection itself and the subsequent processing of acoustic data files. This represents what we see as the first limitation in this field.

A second limitation arises after data have been collected, due to (i) researchers’ understandable focus on specific, often narrowly defined questions, (ii) practical constraints, and (iii) scientific cultural norms that have not encouraged data-sharing. Combined, these factors often lead to a tendency of not publishing or only partially publishing the data collected during research. This lack of data sharing can hinder scientific progress and make it difficult to reproduce research findings (Jenkins et al. 2023; Powers and Hampton 2019; Reichman et al. 2011; Wilkinson et al. 2016); hence, we argue that there is great intrinsic value in publishing fully curated acoustic datasets. If this practice becomes widespread, it would allow scientists to explore a broader range of research questions, improve reproducibility, and facilitate the validation of findings across different studies and populations (Hersh et al. 2023; Powers and Hampton 2019).

In line with this perspective, we present a comprehensive dataset of wild bird songs recorded from a single population of Great Tits (*Parus major*) in Wytham Woods, Oxford, UK. We collected 21,283 hours of continuous recordings across 703 nesting sites over three spring seasons, which resulted in the annotation of over 1,100,000 notes or acoustic units from more than 100,000 songs (see below for definitions of these terms), sung by approximately 400 different male Great Tits. Among these birds, we have detailed information on the identity and life history of 242 individuals, including 50 that were recorded for multiple years. This information includes the time and location of breeding attempts, clutch size, number of fledglings, age of the bird, and basic morphological traits. For birds born in the population (106, or 43% of the total), we also include details such as birthplace, postnatal dispersal distance, mother, and social father.

To complement the song recordings, we have prepared extensive metadata for each of the more than 100,000 songs. This includes details such as the onset and offset times of each note within the song, a song type label, and the time of recording. We also provide the time of the first song during dawn. Finally, we augment the dataset by providing embeddings of each song, which are vector representations derived from a deep metric learning model specifically trained on this dataset. These can be used to identify individuals and in tasks that require similarity judgements.

To the best of our knowledge, this dataset is the largest publicly available collection of bird songs from a single wild population. We hope it will provide valuable insights into a range of scientific questions, including behavioural repeatability and stability, links between vocal performance and reproductive success, the timing of song production, the syntactic organisation of song production, and song learning in the wild.

What follows is a detailed description of the data collection and curation process and the resulting dataset, together with some discussion around potential uses of data presented in this format.

## Data collection

### Study system & fieldwork

Great Tits are small, short-lived birds—average lifespan: 1.9 years—that sing acoustically simple yet highly diverse songs. During the breeding season, from March to June, Great Tit pairs are socially monogamous and defend territories around their nests (Hinde 1952). In Wytham Woods, Oxfordshire, UK (51°46 N, 1°20 W), a population of these birds has been the focus of a long-term study since 1947 (Lack 1964). Wytham Woods is a semi-natural predominantly deciduous woodland that spans an area of approximately 385 hectares and is surrounded by farmland. Most Great Tits in this population breed in nest boxes with known locations (see map in Figure 1), and the majority of individuals are marked with a unique British Trust for Ornithology (BTO) metal leg ring as either nestlings or adults.

**Figure 1.**
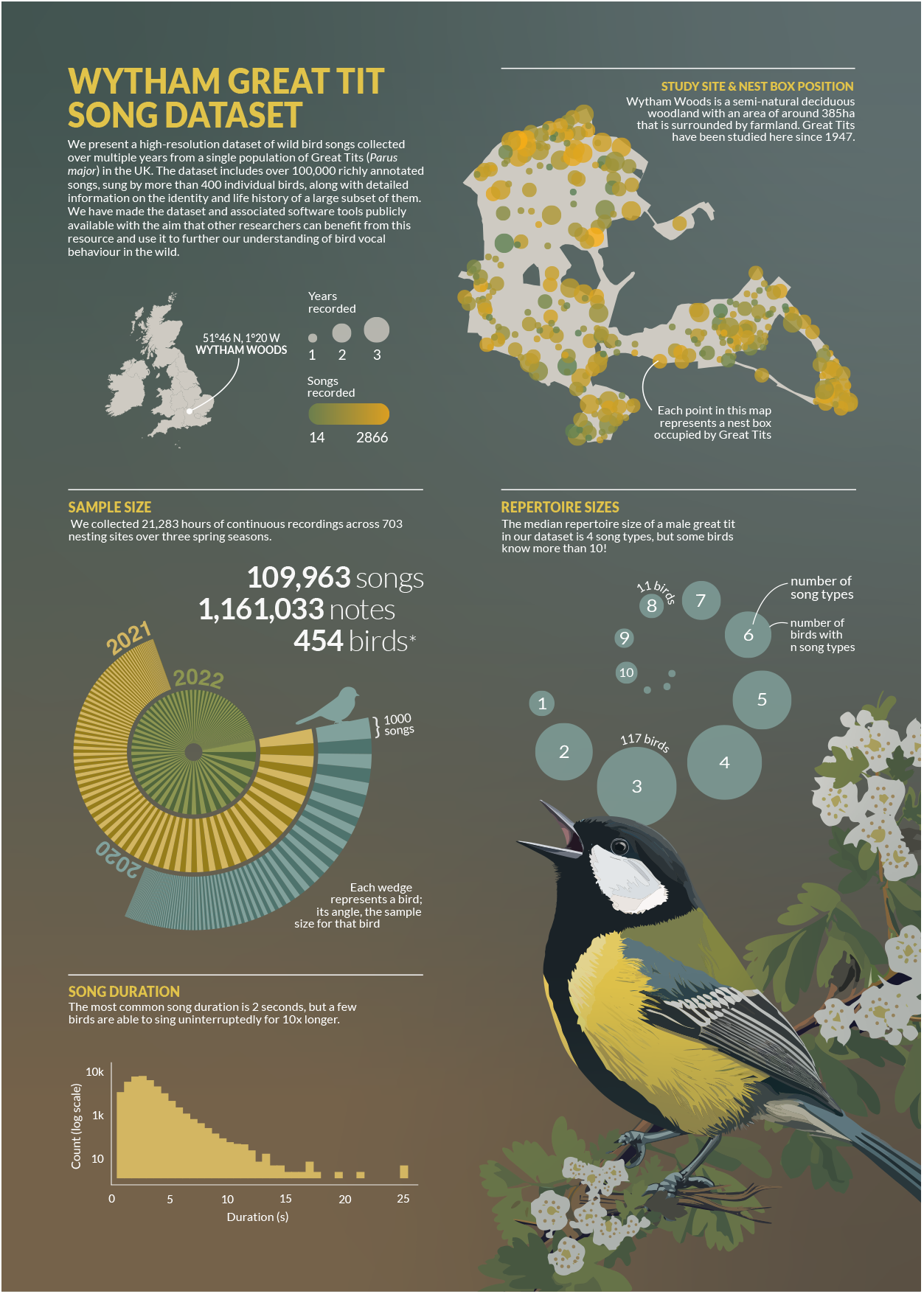
From left to right and top to bottom: Map of the study site and sample locations, total sample sizes for each bird and year, distribution of repertoire sizes, and distribution of song lengths. *The exact number of individual birds is not known exactly.

We collected data from late March to mid-May during the breeding seasons of 2020, 2021, and 2022. Every year, fieldworkers checked each of the 1018 nest boxes at least once a week before and during the egg-laying period, which typically lasts from one to 14 days (Perrins 1965), and recorded the identities of breeding males and females, the dates of clutch initiation and egg hatching, clutch size, and fledgling number and condition under standardised protocols.

To record the vocalisations of male Great Tits we took advantage of their behaviour during the reproductive period, when they engage in continuous singing near their nests at dawn before and during egg laying (Mace 1987). Collectively, this vocal display is referred to as the dawn chorus, and has been demonstrated to yield a reliable estimation of the song repertoire of individuals when recorded in full (Rivera-Gutierrez et al. 2012; Van Duyse et al. 2005). As soon as we suspected that a nest box was being used by a pair of Great Tits—based on nest lining materials, egg size if present, or other signs of activity—we deployed an autonomous sound recorder nearby. These recorders were placed either on the trunk of the same tree or on a nearby tree, between 1 and 2 meters above the ground and no more than 5 meters away, depending on tree availability.

All work involving birds was subject to review by the University of Oxford, Department of Zoology, Animal Welfare and Ethical Review Board (approval number: APA/1/5/ZOO/NASPA/Sheldon/TitBreedingEcology). Data collection adhered to local guidelines for the use of animals in research and all birds were caught, tagged, and ringed by BTO licence holders.

### Recording equipment and schedule

We used 60 (30 in 2020) AudioMoth recorders (Hill et al. 2019), which were housed in waterproof, custom-built enclosures. Recording began approximately one hour before sunrise (05:36 – 04:00 UTC during the recording period) and consisted of seven consecutive 60-minute-long recordings with a sample rate of 48 kHz, and a depth of 16-bit. To sample as many birds as possible, we left each recorder in the same location for at least three consecutive days before moving it to a different nest box. We relocated 20 recorders (10 in 2020) every day throughout the recording period.

### Data processing and annotation

We processed and annotated the recordings using custom software and scripts written in Python 3 (van Rossum 1995), using the open-source package pykanto (Merino Recalde 2023b). These are available from github.com/nilomr/great-tit-hits- setup (Merino Recalde 2023a). Figure 2 shows a graphic illustration of the process.

**Figure 2.**
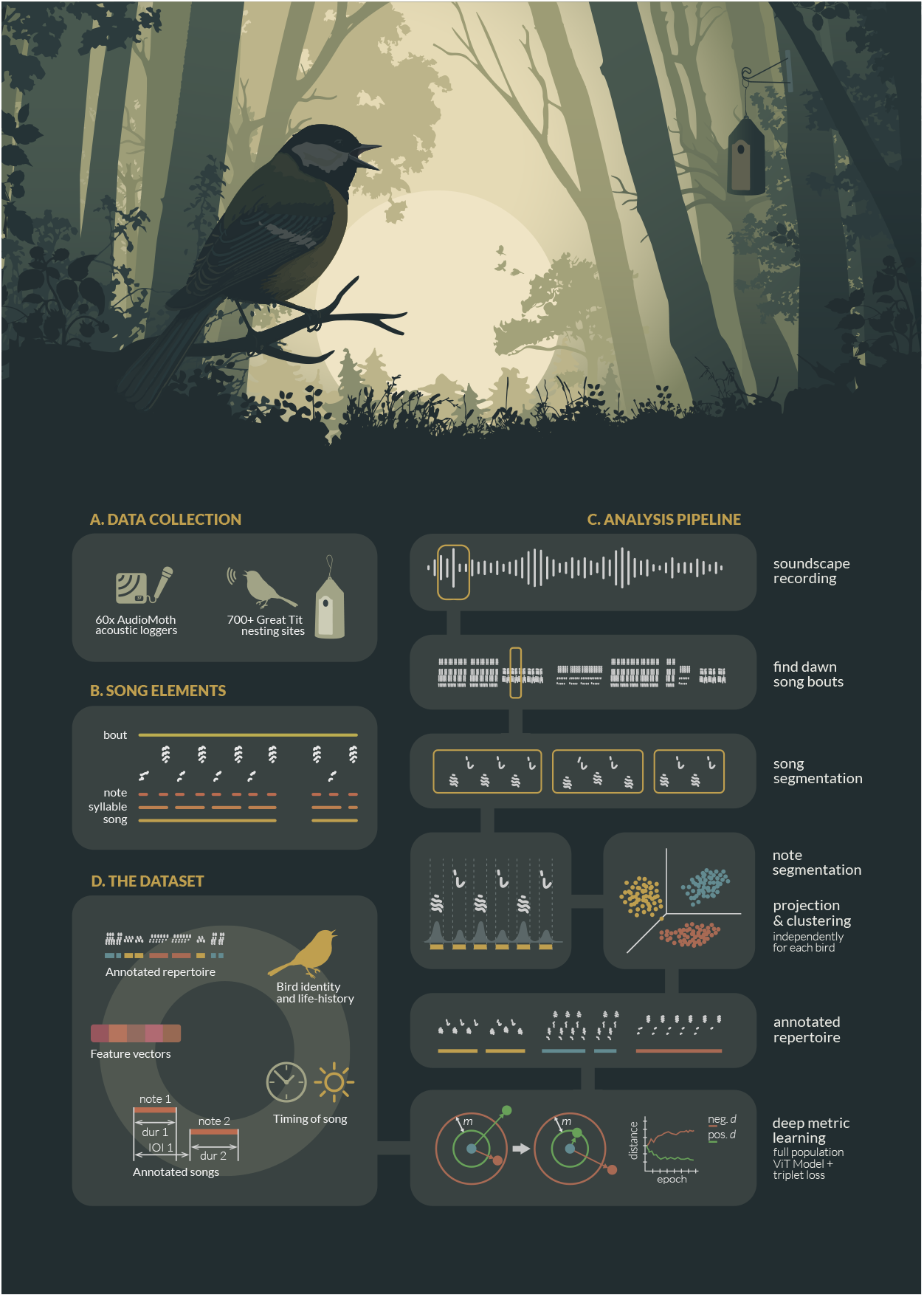
A brief visual summary of the data collection and analysis pipeline used to prepare the Wytham Great Tit Song Dataset. (**A**) Data collection in the field. (**B**) The terminology used to describe the various hierarchical levels at which we can describe Great Tit’s singing. (**C**) Computational pipeline. (**D**) Main outputs included as part of the dataset.

#### A note on terminology

There is no consistent terminology used to describe the various hierarchical levels of a bird’s vocal production. For clarity, we adopt the terminology outlined in (Thompson et al. 1994); see also Figure 2.B for a graphical explanation.

The fundamental temporal unit is referred to as a **note**. Notes are represented by continuous traces on the sound spectrogram and are separated by silences. Moving up the hierarchy, **syllables** are sequences of one or more notes that are always repeated in the same order. Beyond syllables, we have **songs**, which consist of clusters of the same type of syllables punctuated by longer pauses, often in the order of seconds. Lastly, song **bouts** are uninterrupted performances of songs of the same type. Great Tits tend to sing the same song type repeatedly before transitioning to a different type. They continue this pattern until they stop singing altogether, often after having performed their entire song repertoire.

### Song segmentation

We inspected spectrograms for each raw recording and selected songs based on a simple criterion: that its notes were clearly distinct from background noise and other bird vocalisations. We chose entire songs where it was possible; where it was not, we selected the longest contiguous segment possible. This process was carried out manually using the open-source software Sonic Visualiser (Cannam et al. 2010) by drawing boxes bounding songs in the time and frequency domains.

### Assigning song bouts to individuals

Due to the automated recording process, there is a possibility that some of the recorded songs near a particular nest box may not originate from the focal bird. To minimise the chance of false positives, we discarded recordings with more than one vocalizing bird if one was not distinctly louder than the rest during the segmentation process. Additionally, we discarded all songs with a maximum amplitude below −16 dB, calculated as 20 log_10_ 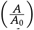, with *A* = 5000and *A*_0_ = 32767 (the maximum value for 16-bit digital audio). This specific threshold was derived from observations indicating that when simultaneous recordings captured neighbouring birds, an amplitude cut-off greater than 4000 consistently differentiated the focal bird from its closest neighbours. It is important to note that these are not calibrated values and are, therefore, relative to the recording equipment and settings we used — as well as other factors like sound directionality and vegetation cover.

### Spectrogramming

For most operations beyond this point, we used normalised, band-passed and log-scaled mel spectrogram representations of each of the songs (sampling rate = 22050, window length = 1024, hop length = 128, mel bins = 224; see the repository nilomr/great-tit-hits-setup for full details on the process).

### Note segmentation

We segmented the resulting song selections into their constituent notes using a custom dynamic threshold algorithm implemented in pykanto (Merino Recalde 2023b), based on the work of Sainburg et al. (2019). Briefly, the algorithm finds minima in the spectral envelope of a spectrogram, which are considered silences; if the length of the signal between these minima exceeds a maximum note duration, a new local minimum is defined that divides the signal into two shorter segments. This is repeated until multiple notes are defined or there are no local minima below a maximum amplitude threshold. Then, segments below a minimum note duration threshold are discarded. To make the algorithm more robust to noise, the spectrogram is subject to morphological transformations and de-echoing before amplitude information is extracted. The de-echoing algorithm implemented in pykanto is based on that in Luscinia (Lachlan 2016), and works by subtracting a delayed version of the spectrogram from itself. We determined minimum and maximum note length ranges by manually segmenting a small, random subset of songs (n = 30).

It should be noted that the automated segmentation process is susceptible to various factors that can influence its accuracy. These include background noise, significant variation in amplitude between notes, attenuation caused by vegetation, changes in the direction of sound production, and even variations in performance where some notes may be much quieter.

As a result, the algorithm may fail to detect or incorrectly delimit certain notes. Despite this, we estimate that approximately 96% of the notes are correctly segmented (0.037 error rate based on a random subset of n = 1048 notes that were checked manually). Still, depending on specific goals, we recommend manual verification of note segmentation if complete accuracy is crucial.

### Song type annotation

We annotated each song type in the dataset using a semi-supervised approach implemented in pykanto. The process involved several steps to ensure accurate classification. First, we generated average unit spectrograms for each song, which provided a concise representation of their temporal and frequency characteristics. Next, we performed non-linear dimensionality reduction using UMAP (McInnes et al. 2018) and a cluster search using HDBSCAN (McInnes et al. 2017) for each bird in the dataset. See (Sainburg and Hedley 2020; Thomas et al. 2021) for similar approaches. This strategy, while useful, often leads to spurious outcomes. For instance, it may separate renditions of the same song type if variation in performance or background noise exists, or if certain song elements are sometimes attenuated. Such variation could be misinterpreted as distinct song types, leading to an overestimation of repertoire size. To address this, we used the interactive app in pykanto to review and split or combine clusters as necessary for each bird. It is worth mentioning that this process would be significantly more challenging in species with highly variable songs: our approach benefited from the Great Tits’ relatively limited repertoires (1 to fewer than 15 song types in our population) and their tendency to produce stable and stereotyped songs.

### Calculating song embeddings

Comparing animal vocalisations poses a significant challenge for researchers. Traditionally, two approaches have been used: visual comparisons of spectrograms and, more recently, measurement of hand-picked acoustic features (Goffinet et al. 2021). However, these methods have limitations when dealing with noise, variations in performance, and changes in syntax (where compositional syntax is not relevant). For instance, if a song with the sequence “tea-cher, tea-cher” is recorded as “cher-tea, cher”, it might be wrongly perceived as highly dissimilar, despite being the same song (see Stowell 2021; Zandberg et al. 2022 for a good overview of these issues). Additionally, these methods often fail to capture high-level features such as the syntactic relationships between notes and other complex spectrotemporal characteristics that cannot be easily characterised by an orthogonal combination of simple acoustic features.

Unfortunately, we cannot rely on the birds’ perceptual judgments (but see recent work in this direction Morfi et al. 2021; Zandberg et al. 2022). In its stead, we adopted a data-driven approach. Our goal was to define a similarity space based on the inherent variation in the data and the only categorical labels that we know are perceptually and behaviorally significant: song types sung by individual birds. Given that Great Tits can recognise each other based on their vocalisations (Lind et al. 1996), we aimed to define a similarity space that not only facilitates similarity-based research but also captures some of the song characteristics that birds themselves might attend to when distinguishing individuals. To do this, we took advantage of recent advances in the fields of deep learning and computer vision.

Below is a simple narrative description of the process. For further details, see the dedicated repository nilomr/open-metric-learning and the OML library (Shabanov 2023).

### Metric Learning and Vision Transformers

Rather than focusing on classification, we aimed to develop semantically meaningful embeddings. To achieve this, we used a Vision Transformer (ViT) model as a feature extractor in a (Euclidean) metric learning task. These models, inspired by the success of transformers in natural language processing applications, process images by splitting them into patches, treating them as tokens similar to words in a natural language (Dosovitskiy et al. 2021; Raghu et al. 2022). In this case, we used the ViT-S/16 architecture (21.7 M parameters), pre-trained on ImageNet using the DINO method (self-distillation with no labels Caron et al. 2021).

### Model Training

During the training phase, we finetuned the ViT model using the Great Tit song dataset. To optimise the performance of the model, we used Triplet loss, a loss function that ensures that the projection of a positive sample, which belongs to the same class as the anchor point, is closer to the anchor’s projection than that of a negative sample, which belongs to a different class, by at least a specified margin (Hermans et al. 2017; Hoffer and Ailon 2018). This loss function enables embedding points of the same class to form clusters without collapsing into a single point, which allows us to also explore differences within song types. While training the model we mined hard triplets—where the negative sample is closer to the anchor than the positive—and used the Adam optimiser with a fixed learning rate of 1 × 10^*−*5^.

### Handling Data Imbalance and Batch Generation

The distribution of song sample sizes per individual in the Great Tit dataset approximately follows a power law, resulting in a significant data imbalance. Although the use of triplet loss already addresses this issue to some extent (Thakur et al. 2019), we adopt a random subsampling strategy where classes with more than 100 samples are reduced to 100 for computational efficiency, classes with fewer than 15 samples are excluded to allow a large enough query/gallery split for validation, and we ensure fair representation during training using a balanced sampler (Hermans et al. 2017). Our batch generation strategy involves uniformly sampling P song types without replacement and sampling K spectrograms for each song type, with replication as necessary. This guarantees that all labels are selected at least once in each epoch.

### Train-Time Data Augmentation

To enhance model robustness and prevent overfitting, we apply various train-time data augmentation techniques (Mumuni and Mumuni 2022; Perez and Wang 2017; Shorten and Khoshgoftaar 2019). These include random cropping in the time domain, dropping out parts of the spectrogram, adding Gaussian and multiplicative noise, equalisation, sharpening, changes to brightness and contrast, blurring, and slight shifting in both time and frequency domains. The latter augmentations are applied within the typical variation in performance observed in the Great Tit vocalisations.

## Results

Our trained model shows very good performance, achieving a mean Average Precision at 5 (mAP@5) of 0.98 and a Cumulative Matching Characteristic at 1 (CMC@1) of 0.98. This indicates that in approximately 98% of the queries made to the similarity space, the returned candidate song type by a bird is the correct one. Errors primarily stemmed from instances where songs of the same type sung by the same bird appeared more than once in the dataset, which happened if a bird survived to the next year. Given that the model was trained on almost 2000 classes, this means that there is enough individual information contained in each song type to distinguish between birds with high confidence, which has important implications for both the study of individuality and population monitoring. See Figure 3 for a visual representation of the embedding space and nearest-neighbour queries.

**Figure 3.**
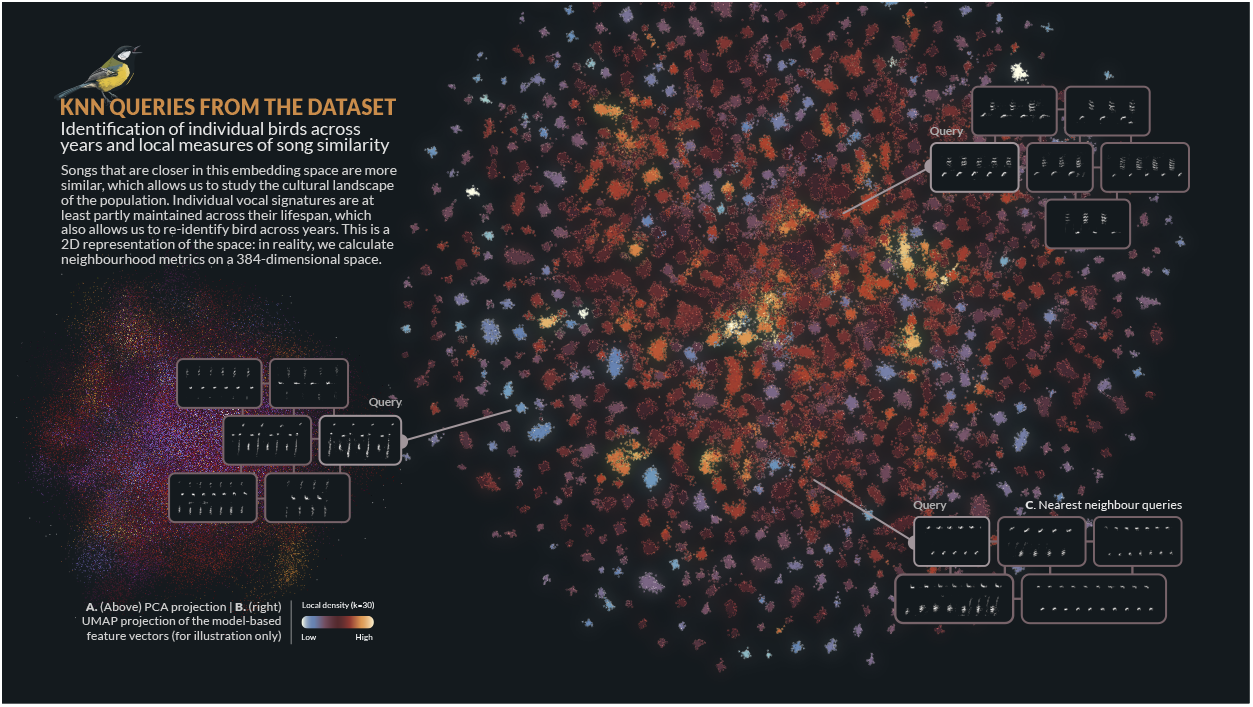
Measuring similarity is a very hard problem, in large part because there is often no objective way to compare the performance of different methods. Here, we took a data-based approach by training a Vision Transformer (ViT) model as a feature extractor in a Euclidean metric learning task. The resulting embedding space allows us to judge if two songs are very similar, and to re-identify birds. (**A**) PCA projection of the feature vectors: two orthogonal linear components do not capture much of the high-level distinguishing features. **B**) This figure shows a UMAP projection of the 384-dimensional vectors for each song in the dataset into 2D, which leads to a fairly arbitrary but useful visualisation where tight clusters of points correspond to song types in the repertoire of individual birds. They are coloured by how densely occupied that region of space is in the high-dimensional space, based on k=30 neighbours from other song types. (**C**) A k-nearest neighbour search returns the closest matches for a query vector (highlighted).

## Data records and description

### Data records

The complete Wytham Great Tit Song Dataset is available at 10.17605/OSF.IO/N8AC9 (Merino Recalde 2023c). All input and output data files use open data formats and are under a CC-BY-4.0 licence. The scripts and software used to create this dataset are available under the MIT licence from GitHub nilomr/great-tit- hits-setup and archived at Zenodo (Merino Recalde 2023a).

Table 1 contains a summary of the files included with the dataset. Detailed data documentation, including variable descriptions, can be found online at nilomr.github.io/great-tit-hits.

**Table 1.** Short description of the files included in the dataset. See the docs for detailed documentation.

|  |
| --- |
| 109,963 raw song files with their corresponding metadata |
| Model-based feature vectors, with 109,963 samples and 384 dimensions |
| A main derived dataset including information on broods, adult bird life-history traits, nesting locations, and acoustic recordings |
| Morphological measurements for birds captured and re-trapped in the study area. Includes information on species, age, sex, weight, and various other traits |
| Information on the location and characteristics of nest boxes in Wytham Woods, along with a map of the site |

### Dataset summary

The dataset provides a comprehensive view of the populations’ natural dawn singing behaviour over three spring seasons. It documents changes in individual performance, the appearance and disappearance of birds—and with them, their songs—and highlights just how much behavioural variation there is along every dimension of what could at first seem a relatively simple trait. Table 2 presents some simple summary statistics, and Figure 1 provides a visual overview of the dataset.

**Table 2.** Dataset sample sizes

|  |  |
| --- | --- |
| Number of segmented notes | 1,161,033 |
| Number of songs | 109,963 |
| Mean repertoire size | 4.24 |
| SD repertoire size | 1.98 |
| Median repertoire size | 4 |
| Number of unique classes | 1,930 |
| Mean class size | 56.98 |
| SD class size | 68.35 |
| Median class size | 31 |
| Number of nest sites recorded | 706 |
| Number of nest sites with data | 454 |
| Number of unique birds with data that were ID'd | 242 |
| Number of times each bird was recorded | 192 (1y), 42 (2y), 8 (3y) |

Even though most birds in the dataset are one or two-year-olds recorded within a single year (which can be attributed to high turnover rates in the population given low annual survival), the dataset includes valuable data on much older individuals, such as a 7-year-old that we recorded in two different years. Among the recorded birds, some display metronome-like regularity in their performance, while others have highly variable or unusual songs, due to learning from allospecific vocalisations, or even issues with their vocal apparatus. You can find some interactive examples at nilomr.github.io/great-tit-hits. The longest song recorded is approximately 20 times longer than the shortest song (and, coincidentally, was sung by one of the largest great tits ever recorded in the Wytham population). The median number of songs per song type and per bird in the dataset is 31, with a significant number of birds having a much larger count, reaching into the thousands. Additionally, the median repertoire size per bird is four distinct song types, although some birds performed as many as 13 distinct song types.

### Known biases and problems

Working with third-party datasets can be challenging, perhaps particularly so in the study of behaviour in natural populations. The familiarity that fieldworkers inevitably develop with the study system and the data is difficult to replace, and, as a result, there is a risk of unintentionally overlooking important sources of bias and variability. We have compiled a list of some key considerations, which, while not exhaustive, can serve as a starting point for identifying and addressing biases when testing hypotheses, estimating parameters, or evaluating findings from the data. These issues can be broadly classified into two groups: those around bird and song type labelling, which can be partially addressed, and those that are inherent in the data or how it was collected.

### Individual and song identification

One factor that can be partially addressed is that the birds recorded in our dataset are not a random subset of the population; they are those that establish territories and begin the breeding process. In turn, birds that are subsequently identified are more likely to be those whose chicks hatch and survive for at least six days, when the first identification attempt is made. This may skew the distribution of certain behaviours within the dataset or lead to endogenous selection bias (Elwert and Winship 2014). One way to quantify the extent to which the subset of identified birds is representative of the entire breeding population would be to compare the distribution of the trait of interest in both groups. See, for example, Kidd et al. (2015), who found that females in nests that fail early in our population are more likely to be immigrant birds breeding in poor-quality areas.

Another issue to consider is that birds may attempt to breed again in the same nest box or elsewhere after a failed attempt. This, coupled with a failure to identify the male associated with those attempts, means that it is conceivable (although likely very rare) that songs from the same bird could appear in the same year twice, leading to pseudoreplication. Similarly, unidentified birds present in the dataset for multiple years could contribute to this problem. One potential way to address these issues is by using song embed-dings for identification based on similarity and assigning dummy IDs to birds believed to be the same individual. At least, this should be modelled to assess the sensitivity of any results to varying degrees of pseudoreplication from this source.

Finally, a few songs might have been mislabelled before model training, as it is not feasible to manually check such a large dataset. However, the model-based embeddings can help identify any mislabeled songs: they will be clear outliers within their respective classes, thanks to the relatively discrete nature of Great Tit repertoires.

### Unequal samples, songs and calls, and female song

As is common in many complex systems, the interaction of the many processes involved in both song production and sampling results in a heavy-tailed frequency distribution of sample sizes. This variation stems from various sources, including characteristics inherent to the study system, such as individual differences in singing activity and temporal fluctuations throughout the spring season. The sampling process introduces further variation, through factors like equipment malfunctions causing small gaps in the data, variation in recording dates relative to peak activity, and the impact of rain and hail on singing activity and recording quality. We cannot assume these processes to be completely independent of each other. Therefore, when analysing song output or repertoire size, it is important to explicitly specify the assumed causal relationship between factors such as individual characteristics, sampling probability, and the outcome measure.

Another important aspect to consider is that, while we have said that the dataset consists of songs, the demarcation between songs and calls is not entirely straightforward. Some vocalisations that would typically be classified as calls, due to their acoustically simpler, shorter, and possibly more stereotyped nature, are actually used as part of the dawn vocal behaviour. These vocalisations are repeated in a manner that creates an impression of functional equivalence to songs. While we have followed criteria similar to other studies (Baker et al. 1986; Fayet et al. 2014; Krebs et al. 1978; Rivera-Gutierrez et al. 2010) to maintain consistency, we believe that this phenomenon warrants further attention. These calls were not segmented and thus are not included in the dataset, but we are happy to provide soundscape recordings to anyone interested in exploring this aspect further.

Finally, although female song in birds has received relatively little historical attention (see Langmore 2020; Odom and Benedict 2018; Riebel et al. 2005 for further discussion), female great tits also sing (see a brief treatment in Gompertz 1961; Hinde 1952). The vast majority of songs in the dataset belong to the dawn song, a behaviour exclusively performed by the male prior to the female leaving the nest (a pattern observed in blue tits as well, as documented by Sierro et al. 2022). Females, on the other hand, vocalise within the nest, but these vocalisations (Gorissen and Eens 2005, 2004) differ from songs and were not typically detectable by our recording devices. Nevertheless, Hinde (1952) suggests that in the absence of males, females may be more inclined to engage in territorial behaviour that involves singing rather than just producing calls. If that is the case, it is possible that our dataset contains some isolated instances of female song.

### Uses and suggestions

The dataset we are presenting contains detailed information about the vocal behaviour and life of wild birds, providing valuable opportunities for investigating a wide range of research questions. In this section, we suggest several research areas that can be explored using this dataset and provide references to relevant studies in the literature.

#### Behavioural repeatability and stability across multiple scales

Researchers can use the dataset to examine the repeatability and stability of song production and song characteristics across different temporal and spatial scales. This includes studying consistency in vocal behaviour within individuals over time and across different contexts, and its links to age (Rivera-Gutierrez et al. 2012; Zipple et al. 2019) and reproductive fitness (Sierro et al. 2023).

#### Links between vocal performance or diversity and reproductive success

Our data can be used to explore the relationships between vocal performance metrics, such as song complexity or vocal diversity, and individual breeding success on a dataset that is much larger than what is typical in the field (Hutfluss et al. 2022; Beecher et al. 2020; Crates et al. 2021; Hiebert et al. 1989; McGregor et al. 1981).

#### Spatial and temporal properties of acoustic communities

The dataset enables investigations into the spatial properties of acoustic communities, including the distribution of singing individuals within a given habitat and across time. This can provide valuable insights into the spatial dynamics of communication networks and acoustic interaction among neighbour birds.

#### Timing and volume of song production

Researchers can use the dataset to analyse the temporal patterns and timing of song production in Great Tits. This might involve studying diurnal variation, seasonal trends, and the influence of environmental factors on the timing and abundance of vocal behaviour. As an example, Figure 4 provides an overview of key temporal shifts in dawn singing behaviour: male birds sing more during the fertile period of the female, and their activity closely tracks advancing sunrise times.

**Figure 4.**
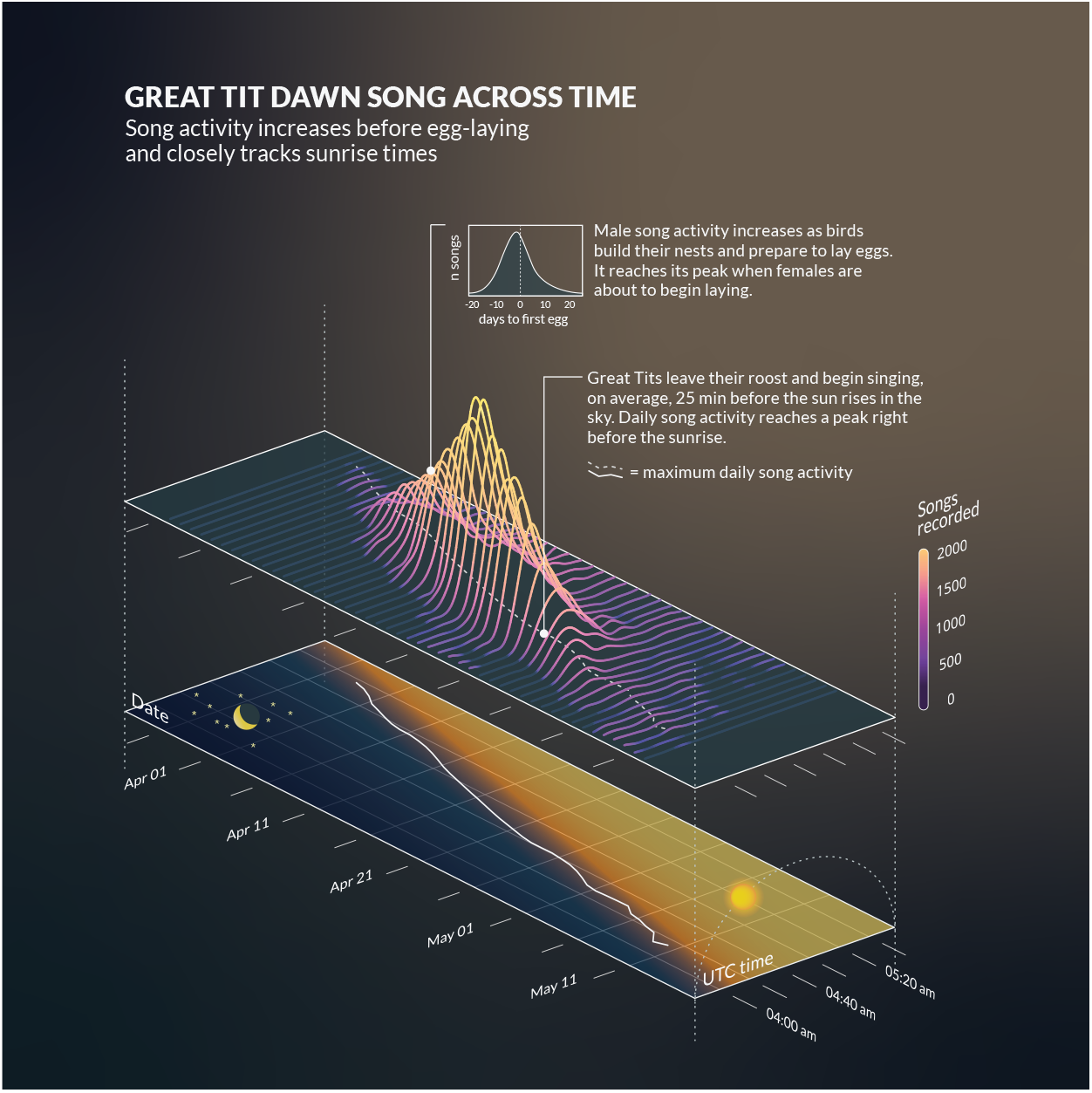
Days get longer as the spring progresses and male Great Tits track the advancing sunrise times with great precision, so that they always begin singing, on average, 25 minutes before the morning breaks. This figure also shows how (z-axis) how song activity peaks alongside egg laying: males sing the most in the morning right before their partner lays the first egg.

#### The syntactic organisation of song production

The dataset captures song activity over entire dawn song periods, across days, and even years for many individuals. This would allow researchers to investigate the set of rules that govern the arrangement of song elements and transitions within the vocal repertoire of wild Great Tits, in terms of short and long-distance dependencies and other properties of their sequential dynamics (Hedley et al. 2018; Lachlan et al. 2010; Sainburg et al. 2019; Searcy et al. 2022).

#### Song learning in the wild

While this dataset does not directly provide evidence of song learning, researchers can use song similarity and proximity in time and space to infer cultural transmission processes. This allows for the exploration of the influence of spatial and social factors on song learning (James et al. 2020; Lachlan and Slater 2003; Nelson and Poe- sel 2014; Peters and Nowicki 2017; Wheelwright et al. 2008).

## Conclusion

With over 1,100,000 annotated notes and acoustic units from more than 100,000 songs, collected over three spring seasons, we hope that this dataset will offer valuable insights into bird vocal behaviour and song culture. The dataset is enriched with detailed metadata such as note onset and offset times, song type labels, and embeddings derived from deep metric learning, as well as identity and life-history information for the birds, which makes it useful for a wide range of research purposes. By sharing this comprehensive dataset, we also aim to help promote data-sharing and scientific collaboration.

## Author contributions

LP, AV, AE and NMR collected the data. NMR created the software and pipeline to plan fieldwork and analyse the data, annotated the dataset with help from AE, built the website, documentation, and visualisations, and wrote the original draft. BCS and EFC provided feedback and supervision throughout the research, and all authors contributed critically to drafts.

## Acknowledgements

We thank all those who have contributed to the long-term nest box study in Wytham Woods and the collection of associated data. This work was supported by a Clarendon-Mary Frances Wagley Graduate Scholarship and an Edward Grey Institute scholarship to Nilo Merino Recalde and made use of the University of Oxford Advanced Research Computing facility (Richards 2015).

## Data and code

The complete Wytham Great Tit Song Dataset and its metadata are available at 10.17605/OSF.IO/N8AC9 (Merino Recalde 2023c). The code, software and pipeline used to create the dataset are available from nilomr/great-tit-hits-setup (Merino Recalde 2023a). The code to train the deep metric learning model can be found at nilomr/open-metric-learning. See the dataset website for documentation and more information.

